# Role of Ori in *Thermococcus barophilus*

**DOI:** 10.1101/2021.04.27.441579

**Authors:** Yann Moalic, Ryan Catchpole, Elodie Leroy, Logan Mc Teer, Valérie Cueff-Gauchard, Johanne Aubé, Yang Lu, Erwan Roussel, Jacques Oberto, Didier Flament, Rémi Dulermo

**Affiliations:** Univ Brest, Ifremer, CNRS, Laboratoire de Microbiologie des Environnements Extrêmes, F-29280 Plouzané, France; Université Paris-Saclay, CEA, CNRS, Institute for Integrative Biology of the Cell (I2BC), 91198, Gif-sur-Yvette, France

**Keywords:** Thermococcus kodakarensis, Ori, RDR, Recombination-dependent replication, DNA replication

## Abstract

The mechanisms underpinning replication of genomic DNA in Archaea have recently been challenged. Species belonging to two different taxonomic orders grow well in the absence of an origin of replication, challenging the role of the replication origin in these organisms. Here, we pursue the investigation of the particular way some archaea manage their DNA replication with *Thermococcus barophilus* and the role of Ori in this Archaea. Surprisingly we discovered that *T. barophilus* uses its Ori all along the growth curve with marked increase at the end of exponential phase. Through gene deletion, we show that Ori utilization requires Cdc6, and that origin deletion results in increased time in lag phase and a moderate decrease of growth rate in mutants. The number of chromosomes are quite similar between both strains during exponential and early stationary phases but differs after 24h of growth where ΔTbOriC has only 6 chromosomes/cell compared to 10 for the reference strain (WT). Following 1hr of growth in fresh media, ΔTbOriC strains contains 3 chromosome copies/cell, whereas the WT contains only 1. We hypothesize that the *T. barophilus* might degrade DNA to obtain energy to start replication and cell division, whereas the ΔTbOriC must maintain more chromosomal copies in order to initiate DNA replication in the absence of an origin or replication. Finally, we analyzed the role of Ori at temperatures above or below the optimal temperature, revealing that Ori is important to start growth at those temperatures, suggesting that replication origins may be involved in stress response.

## Introduction

DNA replication is an essential process for all cells, facilitating the duplication of DNA before cell division. This process begins by the recognition of a specific DNA sequence (Ori) by an initiator protein which promotes opening of the DNA double helix. This role is played by DnaA in Bacteria and Orc1/Cdc6 in Archaea (Leonard and Méchali, 2013). In general, Bacteria have only one Ori sequence (Gao, 2015) whereas up to four have been identified in some Archaea. Indeed, the Archaeal species *Pyrococcus abyssi* and *Nitrosopumilus maritimus* have only one chromosomal replication origin whereas *Sulfolobus acidocaldarius* and *Sulfolobus solfataricus* have three, and *Pyrobaculum calidifontis* has four (Myllykallio et al., 2000; Lundgren el al., 2004; Robinson et al., 2004; Pelve et al., 2012; Pelve et al., 2013).

It was recently shown that origins of replication are not always essential in Archaea; the four origins of *Haloferax volcanii* (DS2 strain) and the single origin of *Thermococcus kodakarensis* can be deleted (Hawkins et al., 2013; Gehring et al., 2017). A slight increase of growth rate was even observed for the multiple Ori depleted strain of *H. volcanii* questioning the role and maintenance of these origins (Hawkins et al., 2013). In the *T. kodakarensis* Ori depleted strain, while growth rates were unaffected, a loss of long-term viability was observed (Gehring et al., 2017). However, not all Archaea tolerate the loss of Ori as was shown for *Haloferax mediterraneii* which must conserve at least one origin of replication to be viable (Yang et al., 2015). Similarly, the Ori-binding protein Cdc6 can be removed in *T. kodakarensis* (Gehring et al., 2017), just as *DnaA* can be removed in cyanobacteria (Ohbayashi et al., 2016); whereas it is not possible to delete all *cdc6*-encoding genes from *H. volcanii* (H26 strain), with some remaining essential (Ludt and Soppa, 2018). Several mechanisms have been proposed and studied for the initiation of replication which are independent of Cdc6/DnaA e.g. rolling circle replication of plasmids by Rep proteins (Ruiz-maso et al. 2015); iSDR (inducible Stable DNA Replication) and cSDR (constitutive Stable DNA Replication) (Kogoma, 1997; Michel and Bernander, 2014). iSDR is a particular form of Recombination-Dependent-Replication (RDR) induced in *E. coli* during the SOS response, initiating chromosomal replication from D-loops (intermediates in homologous recombination). In contrast, cSDR occurs in RNaseH mutants of *E. coli*, where RNA transcripts invade the DNA duplex creating an R-loop sufficient to initiate replication. Both iSDR and cSDR function independently of specific Ori sequences, of protein synthesis, and of DnaA, though both require homologous recombination proteins such as RecA and PriA (Kogoma, 1997; Michel and Bernander, 2014). However, it was proposed that recombination-dependent replication (RDR) could be used in *H. volcanii* since *RadA* became essential in the strain deleted of all four Ori (Hawkins et al., 2013). RDR was first discovered during replication of T4 phage and functions by the use of loops formed after strand invasion to initiate replication of DNA. It was shown that homologous recombination proteins of T4 phage are essential to perform this function (Kreuzer and Brister 2010). The ability of some species to survive without Ori raises many questions, such as the mechanism by which replication occurs in the absence of functional origins, why the origin is maintained when non-essential, and why there is disparity in the essential/non-essential nature of origins between species. Clearly, DNA replication in *Archaea* remains mysterious in many aspects.

In order to better understand the role of Ori in different aspects of Archaeal growth we used the anaerobic and piezophilic Archaeon, *Thermococcus barophilus* MP. This Euryarchaeal species was isolated from hydrothermal vents (Marteinsson et al. 1999) and is genetically tractable (Thiel *et al*. 2014; Birien et al., 2018). We reveal that this archaeon uses its Ori mainly during the end of exponential phase and the beginning of stationary phase and that, despite this clear Ori use, Cdc6 and Ori are non-essential. Analyses of chromosome numbers in WT and Ori-deletion strains show that Ori and ploidy are somehow linked, most markedly during early and late phases of growth. Our results suggest that Ori-utilizing strains resume growth (from stationary phase) more rapidly than Ori-deletion strains, and that this growth resumption is accompanied by a marked decrease in chromosome number. We hypothesize that the polyploidy of *T. barophlius* can be used as an internal energy store to facilitate growth upon nutrient availability. Additionally, we identify temperature-sensitive growth defects of Ori deleted strains, suggesting a role of Ori in stress response.

### NGS read mapping shows activation of OriC in some *Thermococcus*

Recent work has shown that the chromosomal origin of replication (OriC) is non-essential in the archaeon, *T. kodakarensis* (Gehring et al. 2017). Not only can OriC and *Cdc6* be deleted from *T. kodakarensis*, but no evidence of OriC activation is visible when short-read sequencing data is mapped to the chromosome. To assess whether this characteristic is shared by other *Thermococcales*, we mapped Illumina sequencing reads to the genomes of *T. barophilus* and *Thermococcus sp. 5-4*, using *T. kodakarensis* as a control (Fig. 1). For simplicity, the genetic strain Δ*TERMP_00517* used in this paper, it will be called referred to as wild-type (WT). Marker frequency analysis (MFA) shows a weak but not absent use of Ori for *T. kodakarensis* with a slight preference to use Ori during the stationary phase (Fig. 1A). This contrasts with previous work (Gehring et al. 2017) where no Ori use was found, though this is perhaps explainable by the weak signal observed in our analyses. MFA for *T. sp. 5-4* and *T. barophilus* revealed also the use of one Ori at both growth phases but preferentially in stationary phase (Fig. 1B and C). To our knowledge, this is the first time that Ori-dependent replication was found to be used mainly during stationary phase, rather than in exponential phase when DNA replication is expected to mirror cell division.

**Figure 1:**
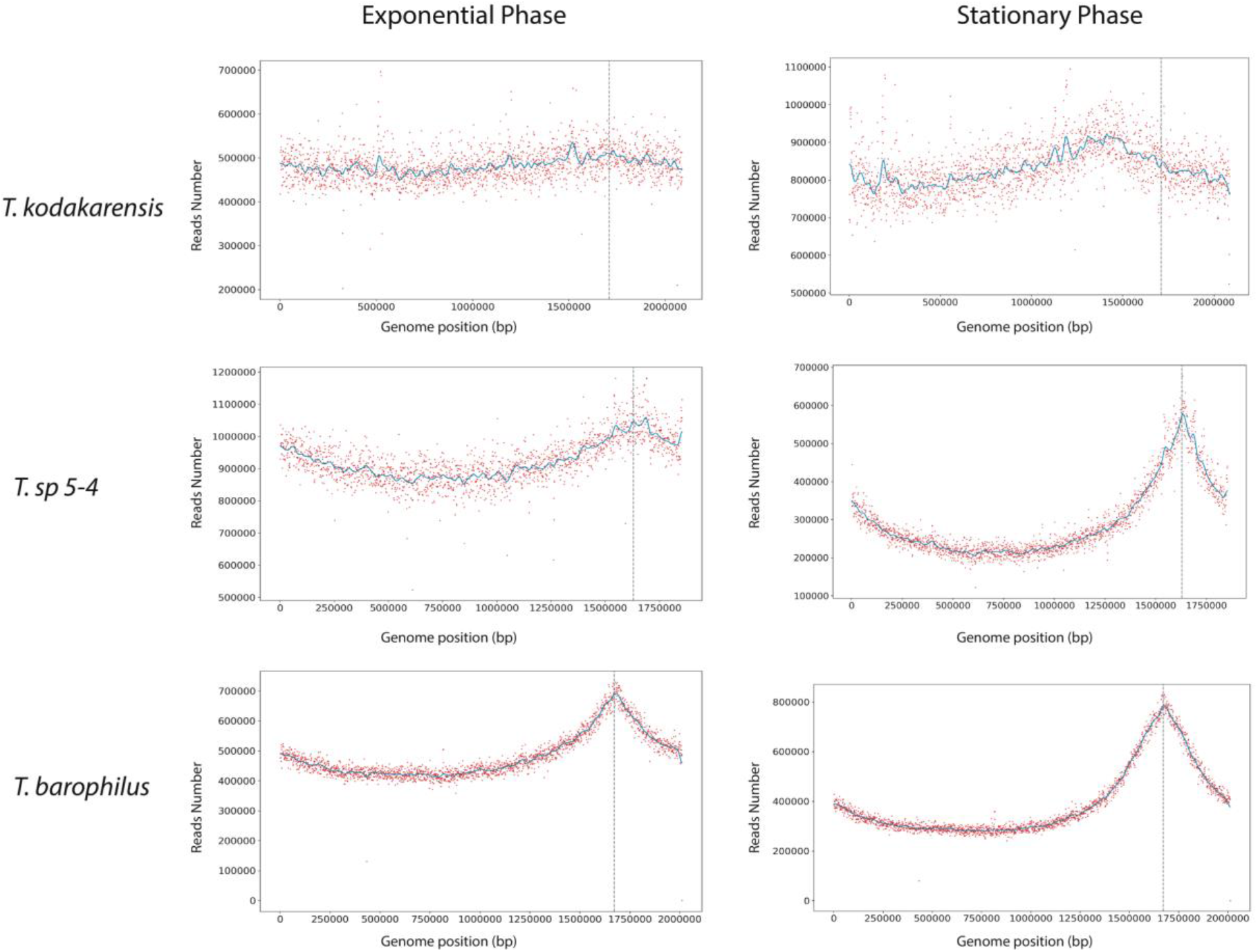
Marker Frequency Analysis of *T. kodakarensis* and *T. sp. 5-4, T. barophilus* genomes. (A) Mapping of Illumina read data to the genomes of *T. kodakarensis* during exponential and stationary phases. (B) Mapping of Illumina read data to the genomes of *T. sp. 5-4* during exponential and stationary phases. (C) Mapping of Illumina read data to the genomes of *T. barophilus* during exponential and stationary phases. The blue lines represent the one dimension Gauss filter. Vertical dotted lines represent canonical Ori position.

### Which Ori is used in *T. kodakarensi*s, *T. sp 5-4* and *T. barophilus*

To localize more precisely the Ori in the 3 species Ori-Finder 2 (Luo et al., 2014) was used to determine the potential Ori sequences. Ori-Finder 2 predicted 3 potentials Ori in *T. kodakarensis*, one localized at 1349201-1349821 (TkOriC1, 2 mini-ORBs; Suppl. Fig. 1A), a second at 1482592-1484232 (TkOriC2, 6 mini-ORBs, Suppl. Fig. 1B) and a third localized at 1711251-1712157 (TkOriC3, 22 mini-ORBs, Suppl. Fig. 1C) adjacent to the *cdc6* gene, corresponding to the Ori found by Matsunaga et al. (2007). *Cdc6* is involved in replication initiation and often found adjacent to the Ori of *Thermococcales* (Ojha and Swati, 2010; Cossu et al. 2015). The first and/or second predicted Ori could be that observed to be active in our MFA.

Ori-Finder 2 found 3 potential Ori for *T. sp 5-4*, at position 1417313-1419615 (5 mini-ORBs, Suppl. Fig. 1D), 1629851-1630129 and 1631511-1632186 (4 and 16 mini-ORBs, Suppl. Fig. 1E). The last two are close to *cdc6* gene. These both positions likely correspond to the active Ori as their coordinates are consistent with the position of the peak observed in MFA (Fig. 1B).

In *T. barophilus*, the software predicted two potential origins of replication, at positions 1333666-1334901 (3 mini-ORBs, Suppl. Fig. 1F) and 1672620-1673707 (20 mini-ORBs, Suppl. Fig. 1G). The latter position (TbOriC) is close to the gene encoding Cdc6 (TbCdc6). Moreover, this position likely corresponds to the active Ori as the coordinates of TbOriC are consistent with the position of the peak observed in MFA (Fig. 1C).

Our data show that Ori prediction are still complicated in *Thermococcus* and that Ori can be found sometimes far from *cdc6* gene. All of this suggest that more effort should be done to understand how replication initiates in *Thermococcales*.

### *T. barophilus* Ori and *cdc6* deletion mutants are viable

It has been shown previously that the origin of replication is not essential for viability in some *Archaea* e.g. *Haloferax volcanii* (Hawkins et al., 2013) and more recently in *T. kodakarensis* (Gehring et al., 2017). Accordingly, TkCdc6 is also not necessary for viability in *T. kodakarensis* (Gehring et al., 2017). In apparent conflict with these results, it was shown that *Haloferax mediterraneii* must conserve at least one origin of replication to be viable (Yang et al., 2015). To determine whether the identified origin of replication and *Tbcdc6* are dispensable in *T. barophilus*, TbOriC and *Tbcdc6* were deleted. Interestingly, both could be deleted, without a strong impact on the growth rate of the strain (Fig. 2A). Sequencing of the mutants shows that they do not contain copies of TbOriC and *TbCdc6*. The doubling time did increase around 18% (75 min for Δ*TERMP*_*00517* to 89 min for Δ*cdc6* and Δ*OriC1*) and a longer lag time was observed (around 6h compared with 3h for WT). In addition, MFA showed that both the TbOriC and *TbCdc6* mutant strains no longer use a detectable Ori (Fig. 2B-E), indicating that TbOriC is the sole Ori used in *T. barophilus* and that it is activated by *Tbcdc6*. The viability of *T. barophilus* in the absence of *Tbcdc6* was similar to that observed for *T. kodakarensis* (Gehring et al. 2017) and *H. volcanii* (Hawkins et al. 2013), and suggests an alternative pathway to initiate DNA replication. This alternative pathway might be mainly observed during the early exponential phase in WT (when TbOriC is weakly used) and at all sample points for ΔTbOriC.

**Figure 2:**
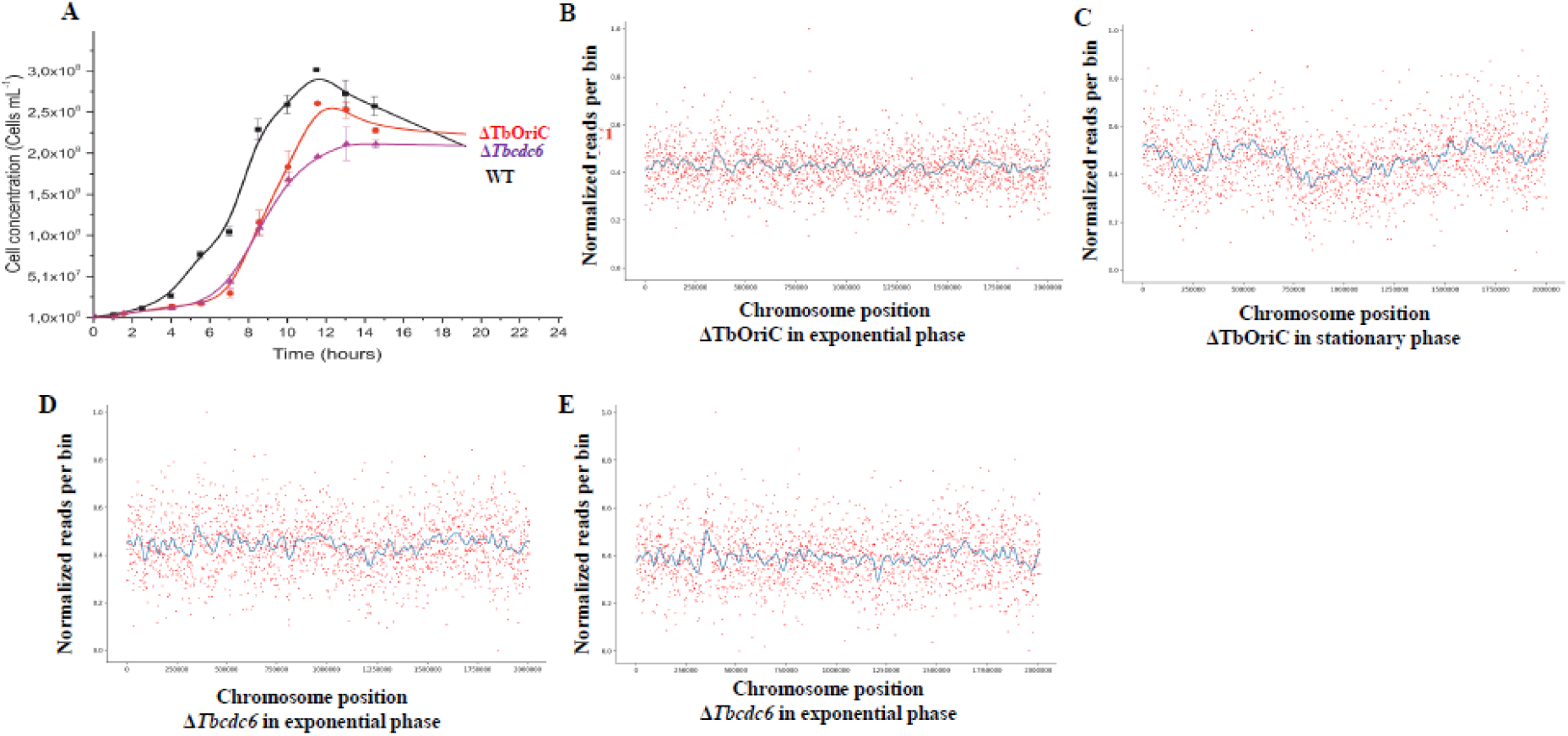
Phenotypes associated with the deletion of *Tbcdc6* and TbOriC. (A) Growth curve of WT, ΔTbOriC and Δ*Tbcdc6* strains. The data presented here are the average of three independent experiments with error bars representing standard deviation. (B and C) MFA of *T. barophilus* strains in the absence of TbOriC (B and C) and *Tbcdc6* (D and E) in exponential (B and D) and stationary phase showing loss of origin usage. The blue lines represent the one dimension Gauss filter.

### Link between Ori utilization and chromosome copy number

To understand how the Ori is activated during growth, we performed MFA on genomic DNA isolated at different phases of growth from both WT and ΔTbOriC strains (Fig. 3, 4A and suppl. Fig. 2). These analyses revealed that WT increasingly utilizes its origin between the middle of the exponential phase to the stationary phase (Fig. 3 and 4B). Interestingly, following transfer to fresh media, we observe a loss of the sharp Ori-utilization peak, with that observed at 4h representing only one fourth of the Ori-dependent replication seen at 1h (Fig. 4B and Suppl. Fig. 2A). Moreover, the width of the Ori peak remains almost stable during the exponential phase (4h to 9.33h of growth, Suppl. Fig. 3), suggesting that most of cells increase Ori opening during this time. This is also correlated with the peak shape that increase in height and has the most pointed shape at 9.33h (Fig. 3), suggesting the highest Ori utilization. During stationary phase, the shape of MFA patterns changes, becoming more round and wider, likely as the Ori-initiated replication propagate around the chromosomes. It is difficult to say whether non-Ori-dependent mechanisms of replication (e.g. RDR) are also occurring in these conditions as these give no signal in MFA; however, parallel experiments showed no Ori activation in ΔTbOriC suggesting another replication mechanism must be at play (Fig. 3, 4B and Suppl. Fig. 2B).

**Figure 3:**
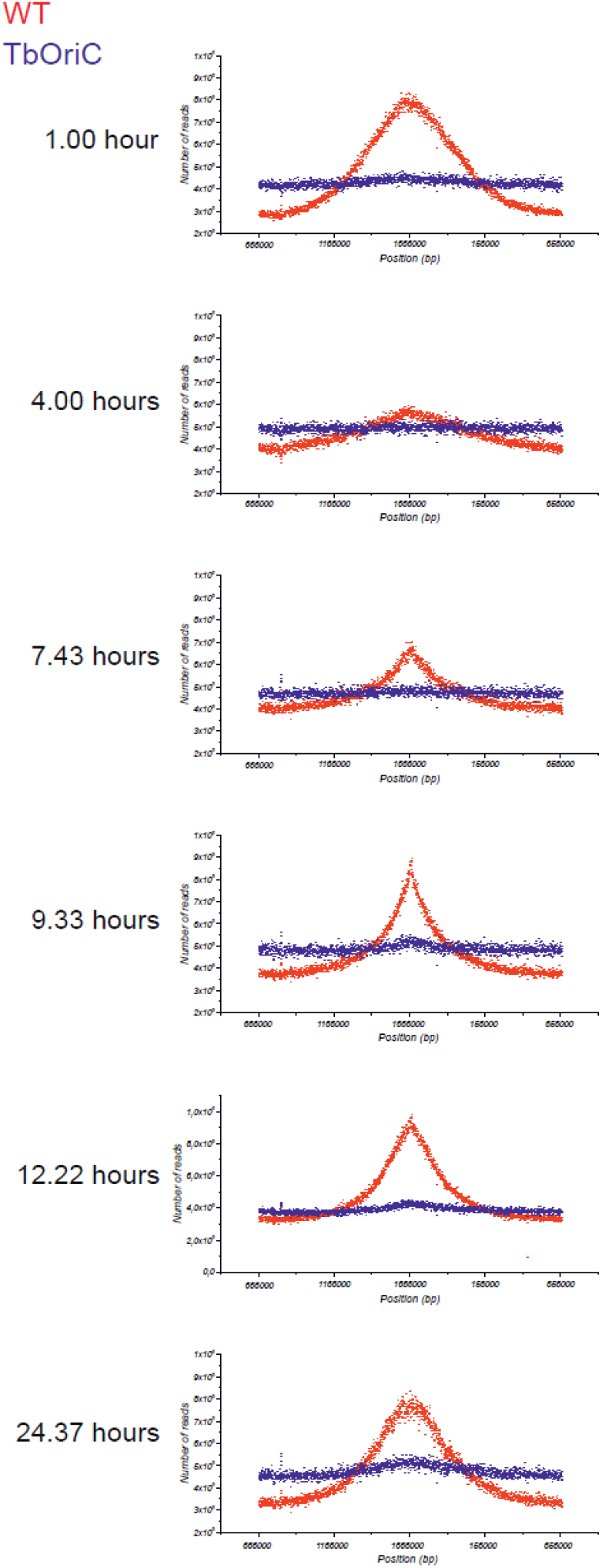
Marker Frequency Analysis of *T. barophilus* WT (red) and ΔTbOriC (blue) strains the long of growth curve.

**Figure 4:**
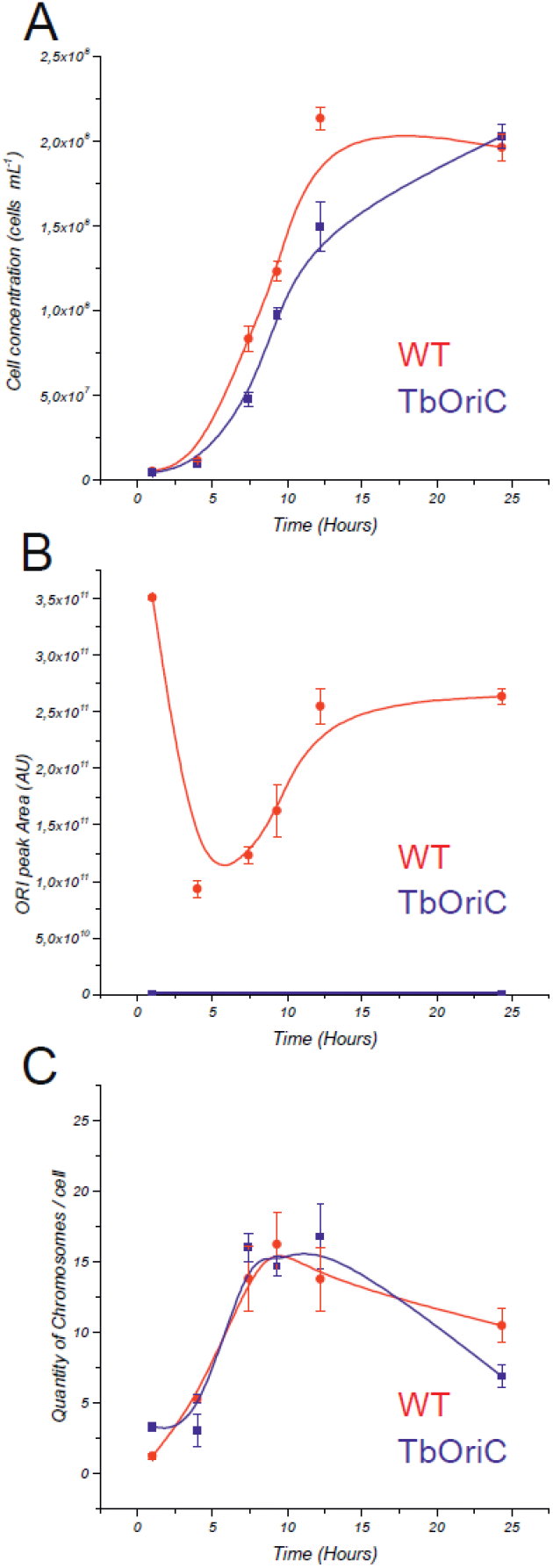
Comparison of Ori peak area and chromosomes number the long of the growth curves of WT and ΔTbOriC. (A) Growth curve, (B) Ori peak area and (C) chromosomes number of WT (red) and ΔTbOriC (blue). The data presented here are the average of three independent experiments with error bars representing standard error.

To understand better how ploidy of *T. barophilus* evolves during growth, qPCR was performed on the *RadA* locus in WT and ΔTbOriC strains (Fig. 4C, Suppl. Fig. 2). Similarly to a previous publication in *Thermococcales* (Spaans et al., 2015), we found that WT contained around 14-16 chromosomal copies during exponential phase and 10-13 during stationary phase (Fig. 4C and Suppl. Fig. 2A). This number was globally similar for ΔTbOriC strain during the exponential phase (around 7h-12h of the growth curve) (Fig. 4C and Suppl. Fig. 2B). However, some discrepancies are observable. Indeed, at 1h of growth, only one chromosome per cell was found in the WT (1.23 ± 0,16) and this correlated with a peak in MFA (Suppl. Fig. 3A) and a short lag phase (Fig. 2). In contrast, around three chromosomes per cell were found in ΔTbOriC (3.29 ± 0,32 chromosomes per cell) suggesting a difference in ploidy regulation or a requirement of polyploidy for growth in the absence of Ori. It is surprising that the number of chromosomes of the WT strain decreased from 10 during stationary phase (10,46 ± 1.22) to one following 1h of incubation in fresh media. We also found that the Ori is necessary to maintain a higher ploidy during stationary phase with Ori deleted strain since it contains only six chromosome per cell at 24h of growth (6,86 ± 0.8), compared with 10 of WT. To better understand the rates of chromosome change in both strains we analyzed the ratios of cells and chromosome number between each time point (Suppl. Fig. 2C). As explained above we could see that at the beginning of the growth, during the 4 first hours, chromosome number per cell increased faster than cell number, showing that at that time WT cells accumulated chromosomes. This contrasts with the ΔTbOri strain where chromosome number stayed constant. During early exponential phase (4h to 7,43h), the ratio of cell numbers increased 7.3 times, faster than the chromosome numbers ratio (2.6 times) in WT, and this without the use of full activated Ori. Again, ΔTbOriC strain behaved differently, with cell number and chromosome number per cell increasing similarly, around 5 times. In middle to late exponential phase (7,43h to 9,33h; 9,33h to 12,22h) we observed a stabilization of chromosome number per cell ratio in both strains whereas, cells number ratios increased slowly. Finally, we could see that between 12,22h to 24,37h, the ratio of chromosome number per cell was below 1, showing a decreasing ploidy. These results suggest that the Ori is important to maintain a higher chromosome number per cell, and perhaps the ability to replicate a single chromosome (Ori-deletion strains never reach n=1). Additionally, a marked reduction in chromosome number seems to correlate with restoration of growth from stationary phase (exit from lag phase).

### Role of TbOriC at sub- and supra-optimal temperatures

*T. barophilus* was described as being able to grow from 48 to 100°C, however, 95°C is the maximum at atmospheric pressure (Martteinsson et al., 1999). To understand the role of Ori in *T. barophilus* under non-optimal conditions, we analyzed growth of WT, ΔTbOriC strains at 65 and 95°C (Fig. 5). As expected, growth rate at those temperatures is lower than at the optimal temperature of 85°C. Generation time for WT is around 162 ± 22 min at 95°C and 494 ± 16 min at 65°C (Fig. 5), compared to 75 min at 85°C. Surprisingly, growth of ΔTbOriC was strongly affected at both 65 and 95°C, not only during the lag phase (as observed at 85°C) but throughout growth, with generation time increased 1.7 time at 95°C (274 ± 23) and around 1.33 at 65°C (657 ± 77 min). This shows clearly that even if growth was not greatly affected by the deletion of Ori at 85°C, this deletion strongly affected growth at 65 and 95°C. The dispensibility of Ori at optimal temperature in *H. volcanii, T. kodakarensis* or in *T. barophilus*, might suggest variation in environmental conditions to be a reason explaining maintenance of Ori and Cdc6 in *T. barophilus*. It remains to be tested whether similar growth defects arise in other species.

**Figure 5:**
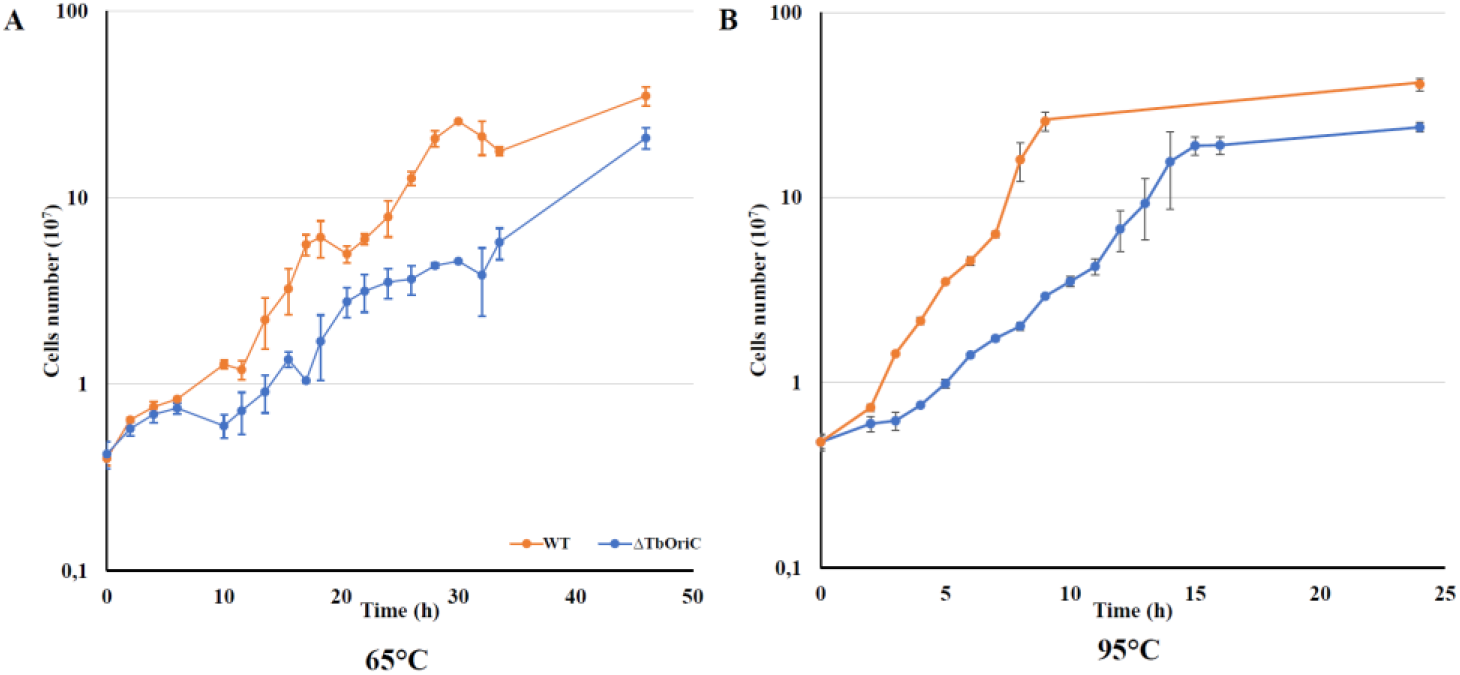
Growth curves at sub and supra-optimal temperature of WT and ΔTbOriC. (A) Growth at 65°C. (B) Growth at 95°C.The data presented here are the average of three independent experiments with error bars representing standard deviation.

## Discussion

Here, we analyzed the role of Ori in *T. barophilus* and found that, in contrast to that known for *T. kodakarensis* (Gehring et al. 2017), a chromosomal origin of replication is used for DNA replication. Surprisingly, this Ori was active mainly during late exponential to early stationary phase. Additionally, we were able to identify subtle Ori use in *T. kodakarensis*, suggesting Ori use may be a regulated process in *Thermococcales* occurring under specific growth conditions.

We were surprised to see that *T. barophilus* use one fourth to one half of the maximal Ori utilization during most of exponential phase growth (results are summarized in Table 1), raising the question about the replication mode of DNA during this phase. Moreover, the functional Ori of *T. barophilus* was dispensable, along with the associated replication initiator, Cdc6.

**Table 1:**
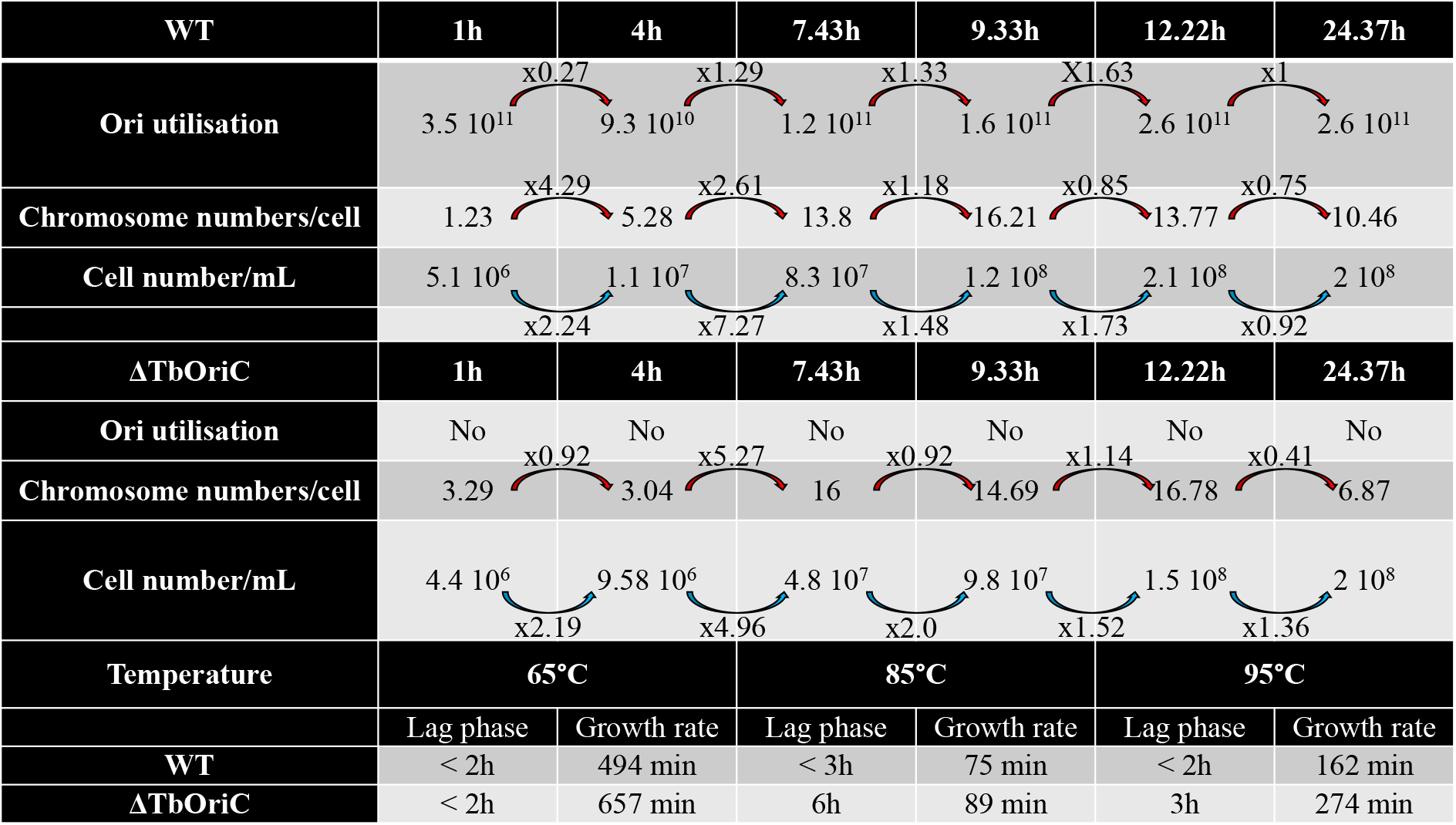
Summary of results

We show that the presence of a functional Ori correlates with reduced lag phase, increased chromosome number per cell, and an improved ability to grow at different temperatures, especially at sub- and supra-optimal temperatures (Table 1). At optimal growth temperature, WT strains appear to rapidly degrade existing chromosomes, decreasing ploidy from around 10/cell in stationary phase to 1/cell following 1h in fresh media. While the cause and significance of this is unknown, it is easy to speculate that such a liberation of resources might facilitate rapid restoration of growth from the stationary phase state. In contrast, cells deleted of the Ori (and/or *cdc6*) are unable to achieve such chromosome losses, harboring only 6-7 chromosomes during stationary phase, and 3 following introduction to fresh media, perhaps resulting in their elongated lag phase relative to WT. It was unexpected to find that at the beginning of the culture, a described polyploid species contains only 1 chromosome per cell. While this is a transient phase, we believe it significant as it is difficult to imagine how DNA replication can proceed from an n=1 state in the absence of an Ori sequence.

All of our data suggest that Ori is important to reduce lag phase, may be by improving preparation of cell for division using DNA as energy stock. In this regard, catabolic pathways exist in *Thermococcales* from which dCTP and dTTP could easily provide substrates for the pentose phosphate pathway or amino acid biosynthesis, for example (Suppl. Fig. 4). For dATP and dGTP, it may also be possible (Suppl. Fig. 4). To prove this hypothesis, more experiments, especially metabolomics, should be performed.

During early exponential growth phase we found that the WT strain does not appear to replicate DNA through an Ori-dependent mechanism (or at least not solely through such a mechanism), with MFA showing a loss of signal at the Ori locus. Despite this, chromosome number per cell continues to increase, suggesting that a potent Ori-independent pathway (e.g. RDR) is sufficient to rapidly replicate DNA. In contrast, ΔTbOriC cannot use Ori to replicate DNA and thus must rely upon Ori-independent mechanisms. It seems that such mechanisms are not as efficient as Ori-dependent pathways to replicate chromosomes and generate a decrease of growth rate observed in Ori deleted strain, especially at 65 and 95°C and to a lesser extend at 85°C. We can see also in our results that Ori-independent mechanisms are not efficient at maintaining high chromosomal copy number in stationary phase, since ploidy decreases more rapidly than in WT strains. This is in accordance with results found by Gehring et al. (2017) where they observed a decrease of viability of ΔTkOri Δ*Tkcdc6* strain when stationary phase is lengthened.

We hypothesize that *Thermococcales* are capable of replicating chromosomes via [at least] two different mechanisms. Upon introduction to fresh media, chromosomes can be degraded and used as a fast source of nutrients to rapidly restore growth. If such chromosome degradation results in an n=1 state, Ori-dependent pathways are required for chromosome replication (assuming the presence of an Ori and associated cdc6). Once ploidy reaches sufficient levels, Ori-independent pathways can proceed and rapidly replicate DNA. Once nutrient density begins to decrease (late exponential/early stationary phase), more conservative replication is preferred, and Ori-dependent pathways again dominate.

It seems unlikely that Archaea replicate via an iSDR mechanism, as iSDR-dependent cells are unable to form colonies in *E. coli* (Michel and Bernander, 2014). Moreover, RNaseH is intact in the studied *cdc6* or Ori mutants of Archaea (including the published *T. kodakarensis* and *H. volcanii*, and our *T. barophilus* strains), suggesting that an alternative form of DNA replication initiation may exist in Archaea. It is worth noting that ΔOri strains of *T. kodakarensis* and *H. volcanii* require the homologous recombination protein, RadA for survival (though this protein has been implicated in other functions of DNA replication). In near future more experiments should be performed to understand where and how precisely DNA replication initiate in Ori deleted strain. It is thus reasonable to propose that some form of recombination-dependent replication is responsible for chromosome replication in exponential growth phase in *Thermococcales* such as *T. kodakarensis, T. barophilus* or *T. sp. 5-4*, and artificially generated Cdc6/Ori mutants. This is supported by the ploidy of ΔTbOriC strain never decreasing below three; perhaps sufficient to permit homologous recombination to start DNA replication.

While our work helps answer why apparently dispensable Ori are preserved in *Thermococcales*, it opens the door to further questions, such as how ploidy is regulated in Archaea, and how Ori-independent replication proceeds.

## Methods

### Strains, media, and growth conditions

Bacterial and archaeal strains are listed in Table S1. *E. coli* strain DH5α was the general cloning host. Luria-Bertani (LB) broth was used to cultivate *E. coli. Thermococcales* rich medium (TRM) was used to cultivate *T. barophilus* MP, under anaerobic condition and at 85°C or at 65 and 95°C, as described in Zeng *et al*. (2009). Artificial sea water complemented by 0.5 % of yeast extract and 0.5 % of tryptone (ASW-YT) was used to cultivate *T. kodakarensis* and *Thermococcus sp. 5-4*, under anaerobic conditions at 85°C, as shown in Sato et al. (2003). Media were supplemented with the appropriate antibiotics used at the following concentrations: ampicillin 25 μg.mL^-1^ for *E. coli*, simvastatin 2.5 μg.mL^-1^ and 6MP (100 µM) for *T. barophilus*. When necessary elemental or colloidal sulfur (0.1 % final concentration) was added for *Thermococcales*. Plating was performed by addition to the liquid medium of 16 g.L^-1^ of agar for *E. coli* and 10 g.L^-1^ of phytagel for *Thermococcales*.

### Plasmids construction

Primers table was given in supplementary table 2. Deletion of TbOriC and *Tbcdc6* was performed using pRD236 and pRD265. These plasmids were constructed using primers pair 145-OriC2c-UpBamHI/250-OriC2c-FusionRv2, 148-OriC2c-DnKpnI/249-OriC2c-FusionFw2 and 298-DeltaTbCdc6BamHI/299-DeltaTbCdc6Rv, 300-DeltaTbCdc6Fw/301-DeltaTbCdc6KpnI, respectively. Fragments generated by PCR were fused using primers pair 145-OriC2c-UpBamHI/148-OriC2c-DnKpnI and 298-DeltaTbCdc6BamHI/301-DeltaTbCdc6KpnI, respectively. Then, these fusions were inserted into pUPH using *Kpn*I and *Bam*HI restriction sites.

### Transformation methods and strains verification

The transformation of *T. barophilus* were performed as already described in Thiel *et al*. (2014) using 0.2 to 2 µg of plasmid. Verification of the deletion was performed using 7-pGDH-IsceI-Fwnew/8-pGDH-IsceI-Rv to ensure that non-replicative plasmid used to constructed mutant did not stay in the cell and for TbOriC, *Tbcdc6* mutants, primers pair outside the construction, 257-VerifTbOriC2-Fw/258-VerifTbOriC2-Rv and 302-DeltaTbCdc6VerifFw/303-DeltaTbCdc6VerifRv were used, respectively.

### Marker Frequency Analysis

DNA was extracted from cultures of *Thermococcus* species at exponential or stationary phase growth (6h and 16h, respectively for *T. kodakarensis* and *T. sp. 5-4*; 6h and 24h for *T. barophilus*) or at different point during the growth curve for *T. barophilus* using protocols described previously (Thiel et al. 2014). Library preparation and Illumina sequencing was performed at Genoscope, France (for *T. kodakarensis* and *T. sp. 5-4*) and Eurofins, Germany or Novogene, UK (for *T. barophilus*). Read mapping was performed with Bowtie2 (Langmead and Salzberg, 2012). Normalized average number of reads per position was used to estimate relative replication initiation activity. High read counts were statistically treaded as peaks and the analysis of the peak area were performed using Origin 2016 (OriginLab Corporation, Northampton, USA) with the peak analyser tool.

### Chromosome number determination by quantitative real-time PCR

Quantitative real-PCR was performed from the different culture samples taken during the growth in triplicates of the wild strain *T. barophilus* and the mutated strain ΔTbOriC from which the DNA was extracted. New primer set specific to *RadA* gene was designed: 539-RadAqPCRFw2 and 540-RadAqPCRRv2. Primer concentration was optimized to minimize the secondary structure formations and to maximize the reaction efficiency. Q-PCR reactions were performed in a final volume of 25 µl using PerfeCTa SYBR Green SuperMix ROX (Quanta Bioscience) on a CFX96 Touch Real-Time PCR System (Biorad), 1 ng of DNA template and 800 nM primers. 40 cycles were performed including one hot-start polymerase activation cycle (95°C, 10 min) and 40 cycles of denaturation (95°C, 15 s) followed by a coupled hybridization and elongation step (60°C, 1 min). Standard curve was obtained from 10-fold serial dilutions (1000 to 109 copies) of plasmid containing RadA gene from *T. barophilus*. Each reaction was run in triplicates. The quality of qPCR runs was assessed based on melting curves and measured efficiencies; the R of standard curves generated by qPCR and efficiency of the reaction were around 0.999 and 90%, respectively. qPCR results were expressed in number of chromosomes per cells.

### Data access

*T. kodakarensis* and *T. sp 5-4* raw reads are available on NCBI under accession number NC_006624 and NZ_CP021848 respectively. *T. barophilus* raw reads are available in the European Nucleotide Archive under Bioproject accession number PRJEB40197.

## Acknowledgment

This work was supported by Ifremer, the “Laboratoire d’Excellence” LabexMER (ANR-10-LABX-19) and co-funded by a grant from the French government under the program “Investissements d’Avenir”. I want to dedicate this work to my mum, Monique Dulermo who permitted me to become a scientist.

## Author Contributions

YM: sequencing and analysis of MFA for *T. barophilus*, writing, discussion; RC: sequencing and analysis of MFA for *T. Sp. 5-4* and *T. kodakarensis* writing, discussion; YL: discussion; EL: growth curves; LMT: growth curves; JA: Sequencing and analysis of sequences for *T. barophilus*; ER: growth curves; JO: discussion and funding; DF: administrative struggle to organize the night shift needed to acquire growth data, discussion, writting and funding; RD: funding, writing, design experiments and construction of plasmids and strains.

