## Supplementary figures and images for "Role of Ori in *Thermococcus barophilus*"

### Fig S3

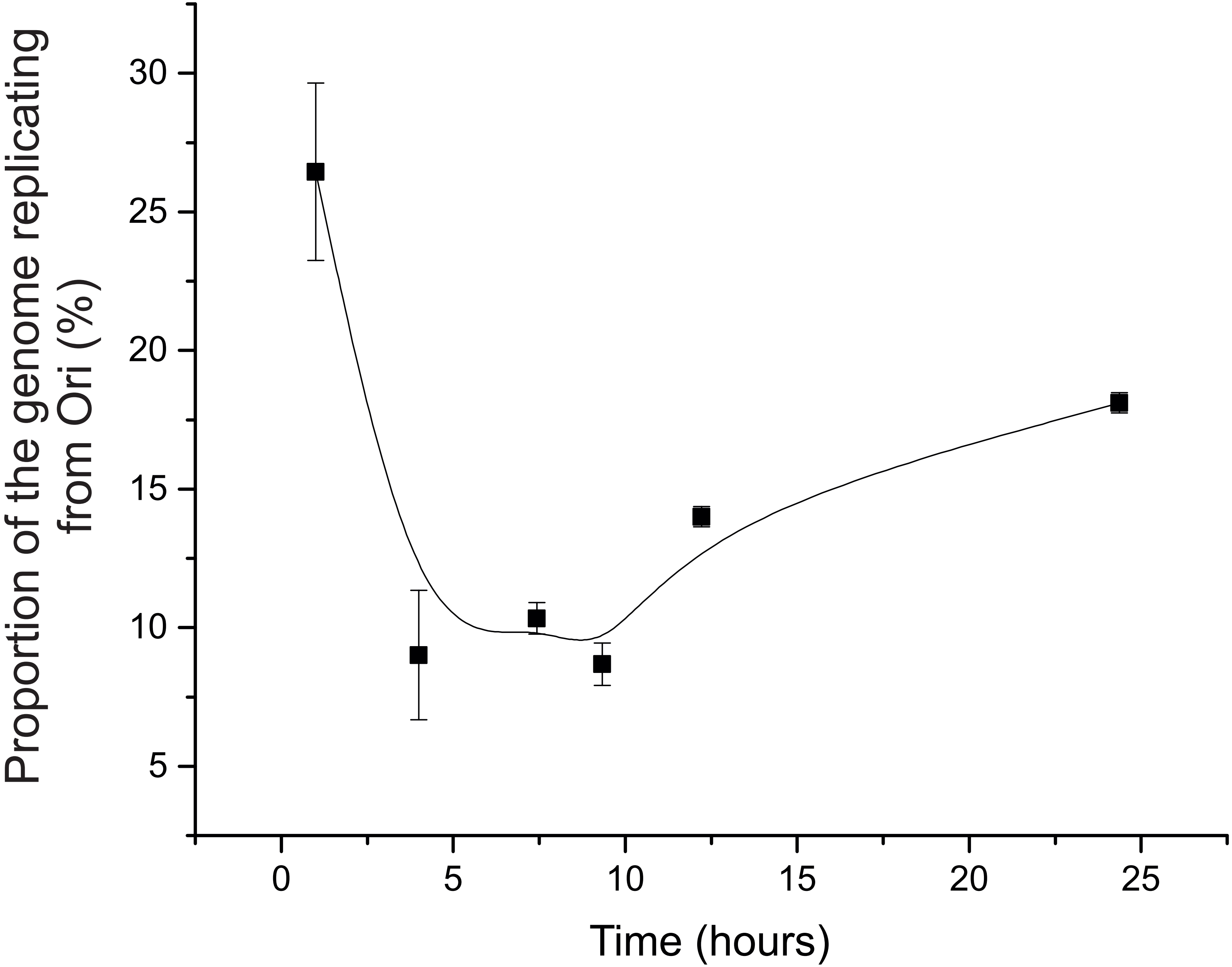

### Fig S4

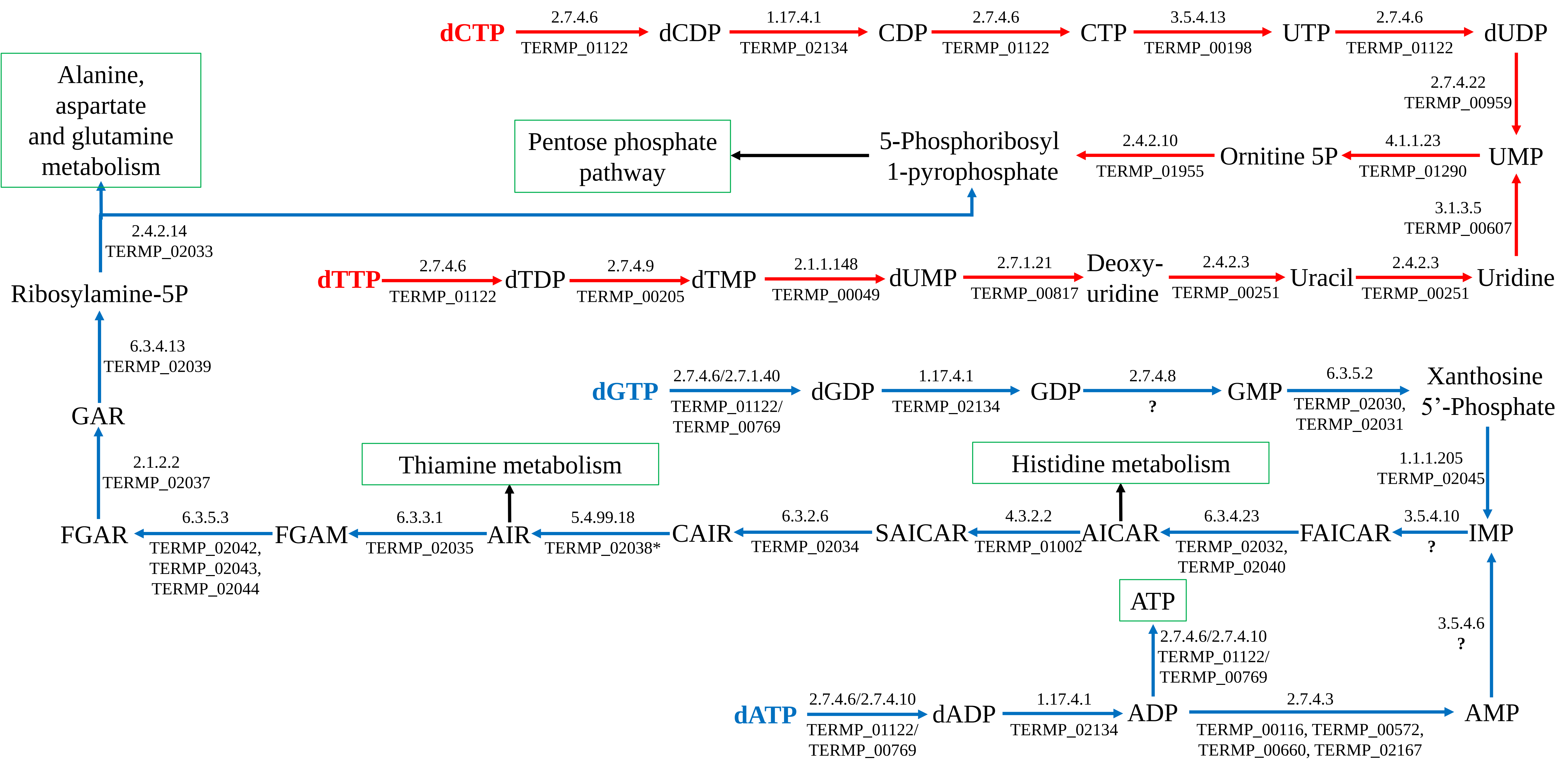

### Figure S1

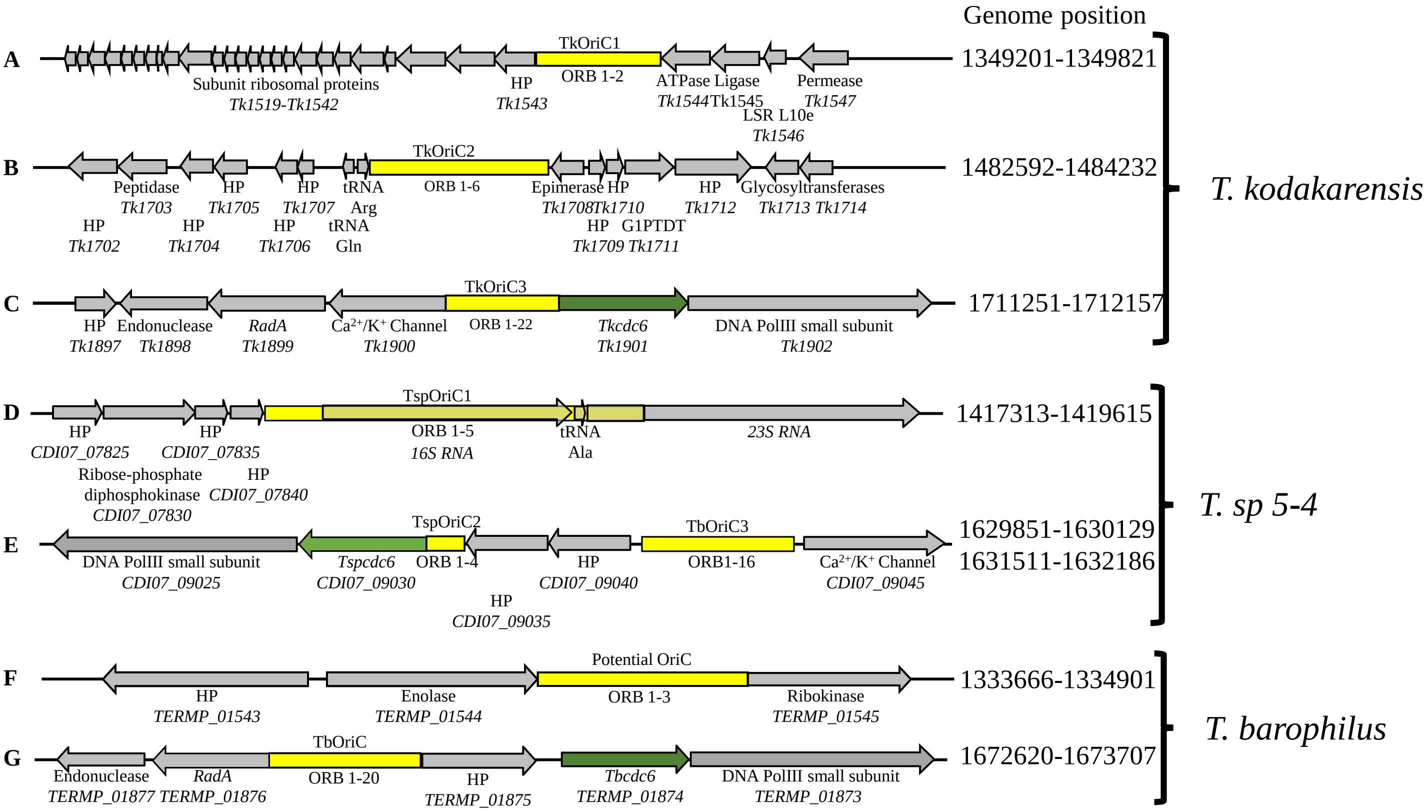

### Figure S2

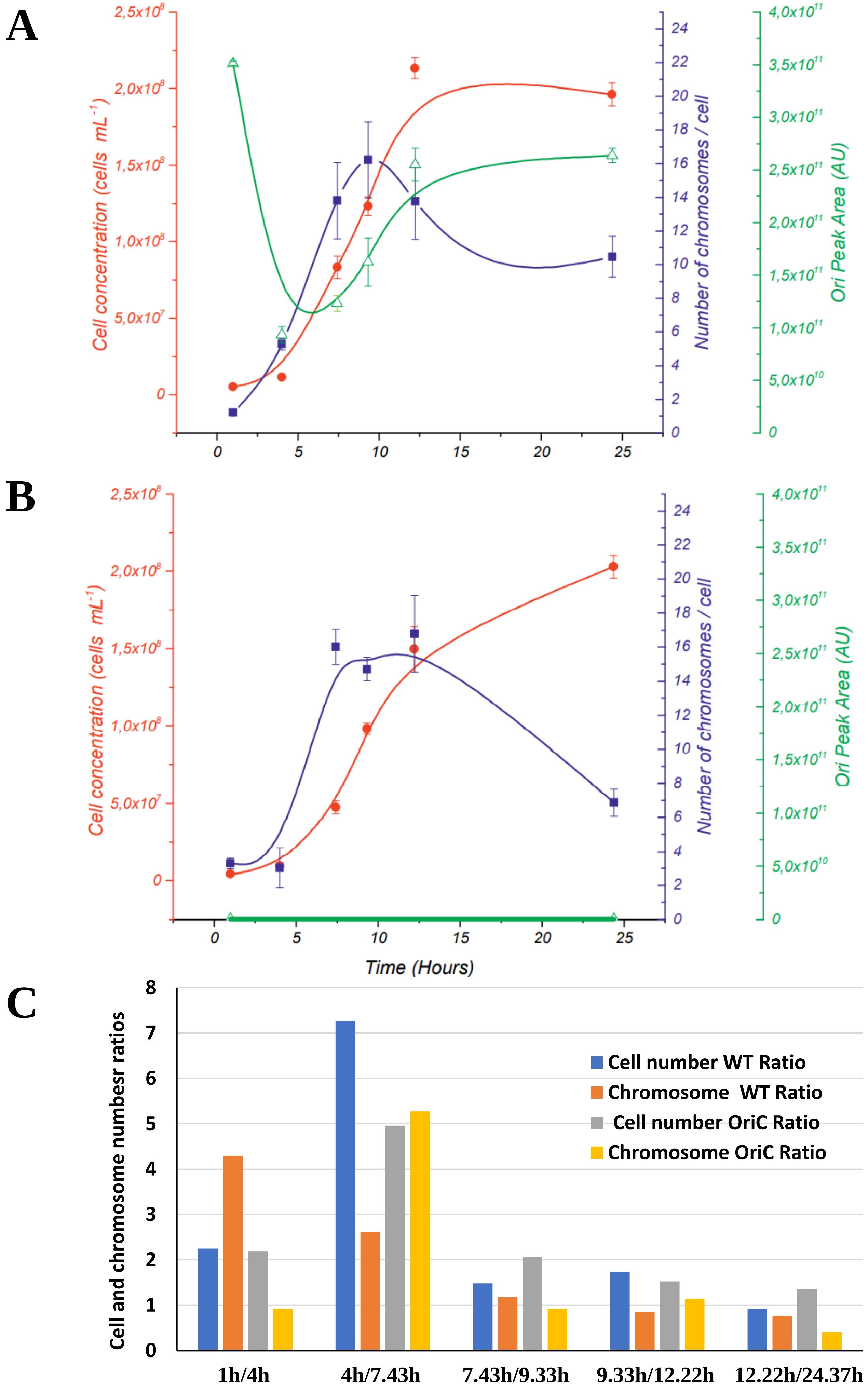
