## Supplementary material for "Role of Ori in *Thermococcus barophilus*": Suppl legends

Supplementary figure 1: Ori-Finder 2 predicted Ori in different *T. kodakarensis*, *T. barophilus* and *T. sp 5-4* and genetic context. (A) TkOriC1, first predicted Ori in *T. kodakarensis* found at position 1349201-1349821. (B) TkOriC2, second predicted Ori in *T. kodakarensis* found at position 1482592-1484232. (C) TkOriC3, corresponding to canonical Ori, is the third predicted Ori in *T. kodakarensis* found at position 1711251-1712157. (D) Potential TbOri, predicted Ori in *T. barophilus* found at positions 1333666-1334901. (E) TbOriC, canonical Ori, at positions 1672620-1673707, close to *TbCdc6* gene. (F) TspOriC1, first predicted Ori in *T. sp 5-4* found at positions 1417313-1419615. Origin underlaps with 23S RNA, tRNA Ala and a part of 16S RNA. (G) TspOriC2 and TspOriC3 second predicted Ori of *T. sp 5-4* found at positions1629851-1630129 and 1631511-1632186, respctively. Ori are represented in yellow and *cdc6* in green. ORB: origin reconition boxes; their numbers, predicted by Ori-Finder 2 are indicated. HP: Hypothetical protein.

Supplementary figure 2: Growth curve, chromosomes number and Ori peak area of WT and ΔTbOriC. (A) Growth curve (red), chromosomes number obtained by qPCR (blue) and Ori peak area (green) of WT. (B) Growth curve (red), chromosomes number (blue) and Ori peak area (green) of ΔTbOriC. The data presented here are the average of three independent experiments with error bars representing standard error. (C) Cell and chromosome numbers per cell ratios between each point.

Supplementary figure 3: Width of Ori peak the long of the WT growth curve. The data presented here are the average of three independent experiments with error bars representing standard error.

Supplementary figure 4: Pyrimidine and purine pathways. Both pathways are represented in red and blue respectively. Green squares represented energy or pentose phosphate pathway. This figure was generated using Kegg website for *T. barophilus* (<https://www.genome.jp/kegg-bin/show_organism?menu_type=pathway_maps&org=tba>). FAIRCAR: 5-Formamido-1-(5-phosphoribosyl)imidazole-4-carboxamide; AICAR: 5-Formamido-1-(5-phosphoribosyl)imidazole-4-carboxamide; CAIR: 5'-Phosphoribosyl-4-carboxy-5-aminoimidazole; AIR: 5-Amino-1-(5-phospho-D-ribosyl)imidazole; FGAM: 5'-Phosphoribosylformylglycinamidine; FGAR: N-Formylglycinamide ribonucleotide; GAR: Glycinamide ribonucleotide. * Sometimes two enzyme are necessary, PurE and PurK but PurK is rare in *Archaea* and PurE could be enough to perform reaction (Brown et al., 2011).
