## Supplementary material for "Role of Ori in *Thermococcus barophilus*": Suppl Table 1

Supplementary Table 1: strains and plasmids

| Strains | Genotype or other relevant characteristics | Source or reference |
| --- | --- | --- |
| ***E. coli*** |  |  |
| DH5α | *Φ80dlacZ*Δ*m15, recA1, endA1, gyrA96, thi-1, hsdR17 (r_k_−, m_k_+), supE44, relA1, deoR,* Δ*(lacZYA-argF)U169* | Thermo Fisher Scientific, Asnières, France |
| ***T. barophilus*** |  |  |
| UBOCC-M-3300 | Δ*TERMP_00517* | Birien *et al*., 2018 |
| RDMP44 | Δ*TERMP_00517* ΔTbOriC | This study |
| RDMP45 | Δ*TERMP_00517* Δ*Tbcdc6* | This study |
| ***T. kodakarensis*** |  |  |
| TS559 | Δ*PyrF*, Δ*TrpE*::P*yrF*, Δ*TK0664*, Δ*TK0149* | Santangelo et al. 2010 |
| ***T. sp. 5-4*** | WT strain | Cossu et al. 2017 |
| **Plasmids** |  |  |
| pUPH | Pop-in Pop-out vector | Birien *et al*., 2018 |
| pRD236 | pUPH + TbOriC2c UpDn | This study |
| pRD265 | pUPH-TbCdc6c UpDn | This study |
