## Supplementary material for "Role of Ori in *Thermococcus barophilus*": Suppl Table 2

Supplementary Table 2: Primers

| Primers name | Sequences | Utilization |
| --- | --- | --- |
| 145-OriC2c-UpBamHI | GCTAGGATCCGGGGTGAATCAATGAGCCTTGC | To delete TbOriC |
| 148-OriC2c-DnKpnI | TCAGGGTACCATTTCCCTGACCCTCCAGTGG |  |
| 249-OriC2c-FusionFw2 | CAAAAGAAGTAAAGTTGATTTTGGACGAATGAATTCCTAAAATTATATTTTAAAGGACAAATGCTAATATTTCTCTGG |  |
| 250-OriC2c-FusionRv2 | CCAGAGAAATATTAGCATTTGTCCTTTAAAATATAATTTTAGGAATTCATTCGTCCAAAATCAACTTTACTTCTTTTG |  |
| 257-VerifTbOriC2-Fw | GGCTGCCTCTCCTTCGGG | To analyze the deletion of TbOriC |
| 258-VerifTbOriC2-Rv | GCAATTCTTTTGGAGTATAGCTATGTCTAAGG |  |
| 298-DeltaTbCdc6BamHI | GCTAGGATCCAACAAGTCATTCAGTGGCTGAGGG | To delete *Tbcdc6* |
| 299-DeltaTbCdc6Rv | CGAGCTCATTTATTAGATCACTGACCCTTCTTCCCTGACCCTCCAGTGGAAACATAGCC |  |
| 300-DeltaTbCdc6Fw | GGCTATGTTTCCACTGGAGGGTCAGGGAAGAAGGGTCAGTGATCTAATAAATGAGCTCG |  |
| 301-DeltaTbCdc6KpnI | TCAGGGTACCTAGTTCTCATAAACCTTGACTACTACCTCTCC |  |
| 302-DeltaTbCdc6VerifFw | ATTTCTCTGGTGATTTCCTGTGGAGG | To analyze the deletion of *Tbcdc6* |
| 303-DeltaTbCdc6VerifRv | CACTAACCTCTGGATTTTCCCGC |  |
| 539-RadAqPCRFw2 | TGCTGTCTCTCTGCTAAAGCTCCC | qPCR primers to test the quantity of chromosomes |
| 540-RadAqPCRRv2 | TGCTGAAAATAGGGGCTTGGATCC |  |
